# Multimodal epigenetic and enhancer network remodeling shape the transcriptional landscape of beige adipocytes

**DOI:** 10.1101/2025.03.28.645896

**Authors:** Sarah Hazell Pickering, Natalia M. Galigniana, Mohamed Abdelhalim, Anita L. Sørensen, Manuela Zucknick, Philippe Collas, Nolwenn Briand

## Abstract

Epigenetic regulation is a key determinant of adipocyte fate and function, conferring phenotypic plasticity to adipose tissue in response to metabolic and thermal challenges. To understand the spatiotemporal regulation of chromatin during the establishment of a beige thermogenic adipocyte phenotype, we analyzed the transcriptomic, epigenetic, and enhancer connectome dynamics during white and beige adipogenesis. Using a machine learning approach, we find that the white-specific transcriptional program is associated with promoter modulations of H3K2ac levels and chromatin accessibility. In contrast, beige-specific mitochondrial gene expression correlates with promoter changes in H3K4me3 levels. Adipocyte beiging is also mediated by a remodeling of the 3D genome involving the recruitment of short range enhancers targeting fatty acid oxidation and thermogenic genes. These increased promoter-enhancer contacts correlate with increased chromatin opening at sites enriched for C/EBP transcription factor motifs. We notably identify the C/EBP transcription factor NFIL3 as differentially bound between white and beige adipocytes at enhancers regulating PDK4, a key metabolic switch promoting fatty acid oxidation. Our results highlight a multimodal, pathway-specific regulation of the transcriptional program underlying the beige adipocyte phenotype.

## Introduction

Adipose tissue plays a central role in whole-body energy homeostasis, supported by the ability of adipocytes to modulate their function and phenotype to adapt to metabolic and thermal challenges. White adipocytes harbor a large lipid droplet *in vivo* and are dedicated to the storage of excess energy in the form of triglycerides, which can be mobilized as free fatty acids to supply energy during fasting periods. In contrast, beige adipocytes display smaller lipid droplets, a high mitochondrial content, and can dissipate energy through thermogenesis, enabled by the expression of uncoupling protein 1 (UCP1) and by the activation of various “futile” cycles (Sharma et al. 2024). Beige adipocytes arise within adipose tissue via *de novo* differentiation and/or from trans-differentiation of white adipocytes in response to environmental cues, such as chronic cold exposure, exercise or treatment with peroxisome proliferator-activated receptor-γ (PPARγ) agonists (Kajimura et al. 2015). This “beiging” process results in whole-body metabolic improvement due to elevated energy expenditure, and increased clearance of triglyceride-rich lipoproteins and glucose from the circulation, making beige adipocytes a potential therapeutic target for metabolic diseases (Bartelt and Heeren 2014).

Epigenetic regulation is a key determinant of adipocyte fate and functional plasticity. Indeed, histone modifications modulate chromatin accessibility, transcription factor (TF) binding, and chromatin folding to control transcriptional activity and gene expression. In mouse models, the identity shift between white and beige adipocytes is accompanied by an epigenetic ‘recoding’ of enhancers (Roh et al. 2018), including acetylation and methylation of histones (Pan et al. 2015; Zha et al. 2015). Although such adaptive mechanisms also likely operate in humans, the epigenetic remodeling and subsequent changes in 3D chromatin organization conferring thermogenic potential to human adipocytes are not fully elucidated.

Here, we characterized the spatiotemporal regulation of chromatin conformation during the establishment of a beige thermogenic adipocyte phenotype. Our integrated, genome-wide, analysis of gene expression in relation to chromatin state, accessibility and 3D conformation highlights multiple levels of regulation mediating beige adipocyte identity.

## Results

### Transcriptional kinetics of white and beige adipogenesis

Human adipose stem cells (ASCs) were differentiated into white or beige adipocytes in the absence or presence of 1 µM rosiglitazone, respectively. Both protocols lead to a similar differentiation efficiency as assessed by Oil Red O staining of neutral lipids at differentiation endpoint (day 15; D15) (**Fig. 1A**), and similar protein expression of PPARG and FAS (**Fig. 1B, C; Supplemental Fig. S1)**. However, beige adipocytes show the expected increase in the protein expression of the beige markers CITED1, UCP1, and of the fatty acid transporter CD36 (**Fig. 1C, D; Supplemental Fig. S1**), and display significantly smaller lipid droplets (**Fig. 1D**). We next performed transcriptomic analysis of white *vs*. beige adipogenesis in a triplicate differentiation time course, starting from a common pre-induction (D0) timepoint. Principal component analysis (PCA) clearly discriminates timepoints (D0, D1, D3, D15) as well as white and beige adipocytes on D15 (**Supplemental Fig. S2**). Indeed, rosiglitazone elicits the upregulation of 1680 genes in beige adipocytes (beige D15 differentially expressed genes; bD15 DEGs), including the beige markers *UCP1*, *CIDEA* and *PEMT*, while 979 are upregulated in white adipocytes (wD15 DEGs) (**Fig. 1E; Table S1**). Strikingly, a large proportion of wD15 DEGs are most expressed on D0 (cluster w2) or at early differentiation timepoints (cluster w1; **Fig. 1F**, left panel). Conversely, bD15 DEGs are almost exclusively late induced (cluster b2, b3; **Fig. 1F**, right panel). Comparative functional analysis across clusters reveals a specific enrichment for extracellular matrix genes among wD15 DEGs (cluster w1, w2), while bD15 DEGs are enriched for genes pertaining to the hallmark “adipogenesis” term, consistent with increased PPARγ activity in rosiglitazone-treated cells (Lehmann et al. 1995) (**Fig. 1G**). Interestingly, genes related to distinct mitochondrial processes are overrepresented in both white and beige D15 DEGs (**Fig. 1G**), suggesting a remodeling of the mitochondrial proteome in beige *vs.* white adipocytes in our system. Supporting this view, white and beige adipocytes display distinct stoichiometries of the mitochondrial respiratory complexes I, II, III and IV, as assessed by Western blotting (**Fig. 1H, I, Supplemental Fig. S3**). Altogether, these results highlight distinct transcriptional kinetics underlying the establishment of white *vs.* beige adipocyte phenotypes.

**Figure 1.**
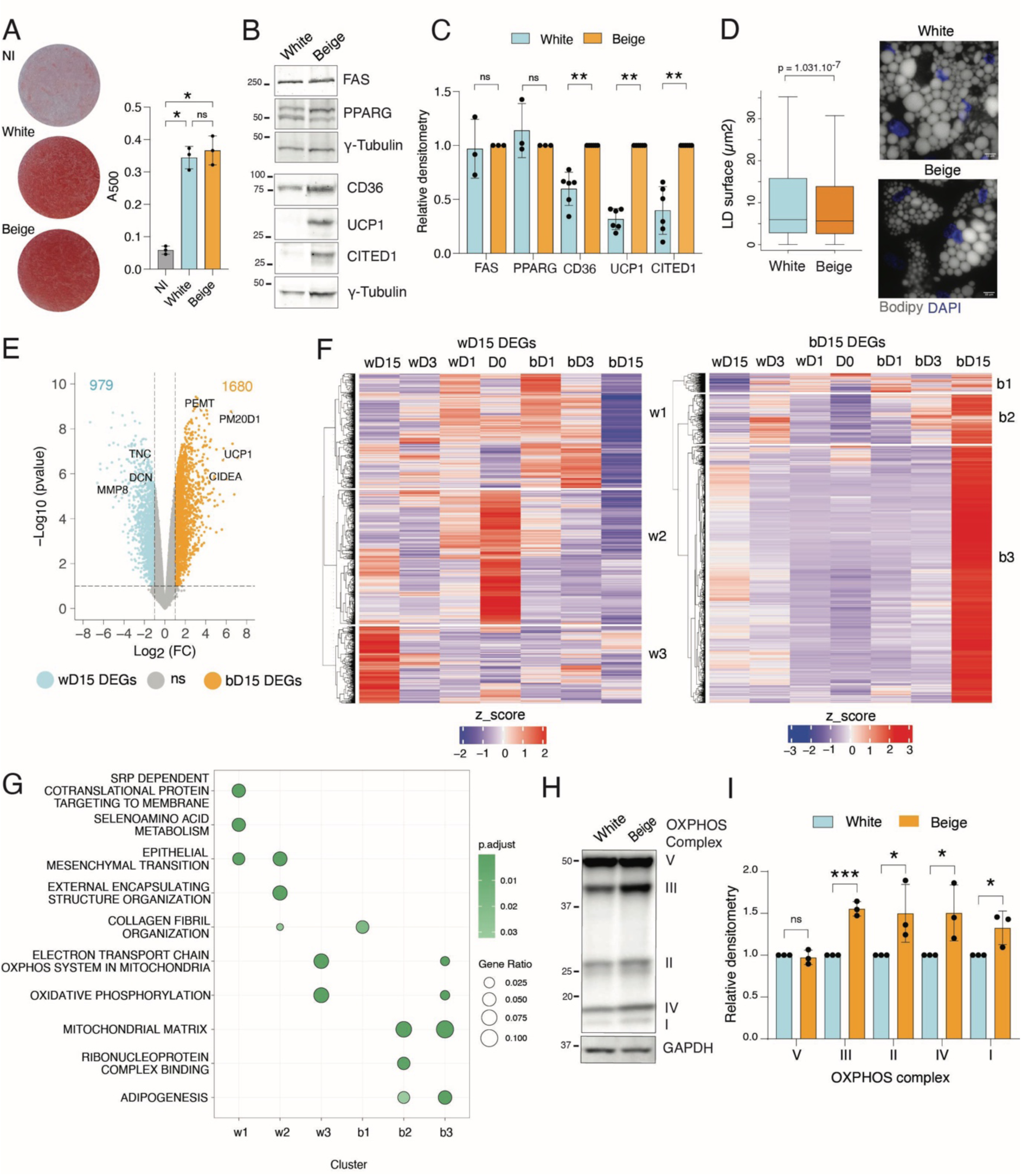
Characterization of white and beige differentiated adipocytes. **A** Representative Oil Red O staining (left) and staining quantification (right) of non-induced (NI) ASCs and differentiated (D15) white and beige adipocytes (mean ± SD; * p < 0.05, ns non-significant, one-way ANOVA with Holm-Šídák’s multiple comparisons test; n = 3 independent differentiations). **B** Western blot analysis and **C** quantification of FAS, PPARG, CD36, UCP1 and CITED1 protein levels in differentiated (D15) white and beige adipocytes (mean fold difference ± SD; ** p < 0.01, *** p < 0.0001, ns non-significant, Wilcoxon signed ranked test; n ≥ 3 independent differentiations). γ-Tubulin is shown as a loading control. **D** Lipid droplet area analysis (Wilcoxon ranked sum test; n = 3 independent differentiations) (left panel) from bodipy staining in differentiated (D15) white and beige adipocytes (right panel); Scale bar: 10 µm. **E** Volcano plot of differential gene expression in differentiated (D15) white and beige adipocytes (p value < 0.01, |log2 fold-change | > 1; ns non-significant). **F** Hierarchical clustering of genes overexpressed in D15 white (wD15 DEGs; left panel) or beige (bD15 DEGs; right panel) adipocytes, scaled across the differentiation time-course. **G** Comparative overrepresentation analysis of wD15 and bD15 DEGs clusters from **F** using GO, Reactome and Hallmark gene sets from MSigDb. **H** Western blot analysis and **I** quantification of OXPHOS complex proteins levels in differentiated (D15) white and beige adipocytes (mean fold difference ± SD; *p < 0.05, ***p < 0.001, ns non-significant, Wilcoxon signed ranked test; n = 3 independent differentiations). GAPDH is shown as a loading control.

### Differential chromatin opening at promoters and enhancers in mature beige adipocytes

To determine to what extent changes in chromatin accessibility contribute to the beige adipocyte phenotype, we performed ATAC-seq during the white and beige differentiation time courses. PCA of ATAC read counts discriminates each timepoint and shows a good reproducibility between replicates (**Supplemental Fig. S4A**), allowing us to call differentially accessible regions (DARs). Overall, the numbers of ATAC peaks and peak coverage are similar between samples (**Supplemental Table S2**). However, induction of adipogenesis elicits tens of thousands of DARs (p < 0.01) resulting in a global increase in accessibility by D15 in both lineages (**Supplemental Fig. S4B**). Beige *vs.* white DARs only emerge on D15, with beige adipogenesis enhancing accessibility in 24,637 regions and reducing accessibility in 17,577 regions (**Fig. 2A)**. D15 DARs are enriched in intronic and distal intergenic regions, suggesting large changes in enhancer accessibility between white and beige adipocytes, with beige DARs showing additional enrichment at proximal promoters compared to white DARs (**Fig. 2B**). In agreement, the vast majority of beige and white DARs annotates to enhancer elements, defined in a chromatin state model learned from our ChIP-seq data (**Fig. 2C**; **Supplemental Fig. S5**), and 20% of beige DAR coverage annotate to TSSs (*vs.* 7% for white DARs). Thus, beige adipogenesis results in increased chromatin accessibility at both enhancer and promoter regions.

**Figure 2.**
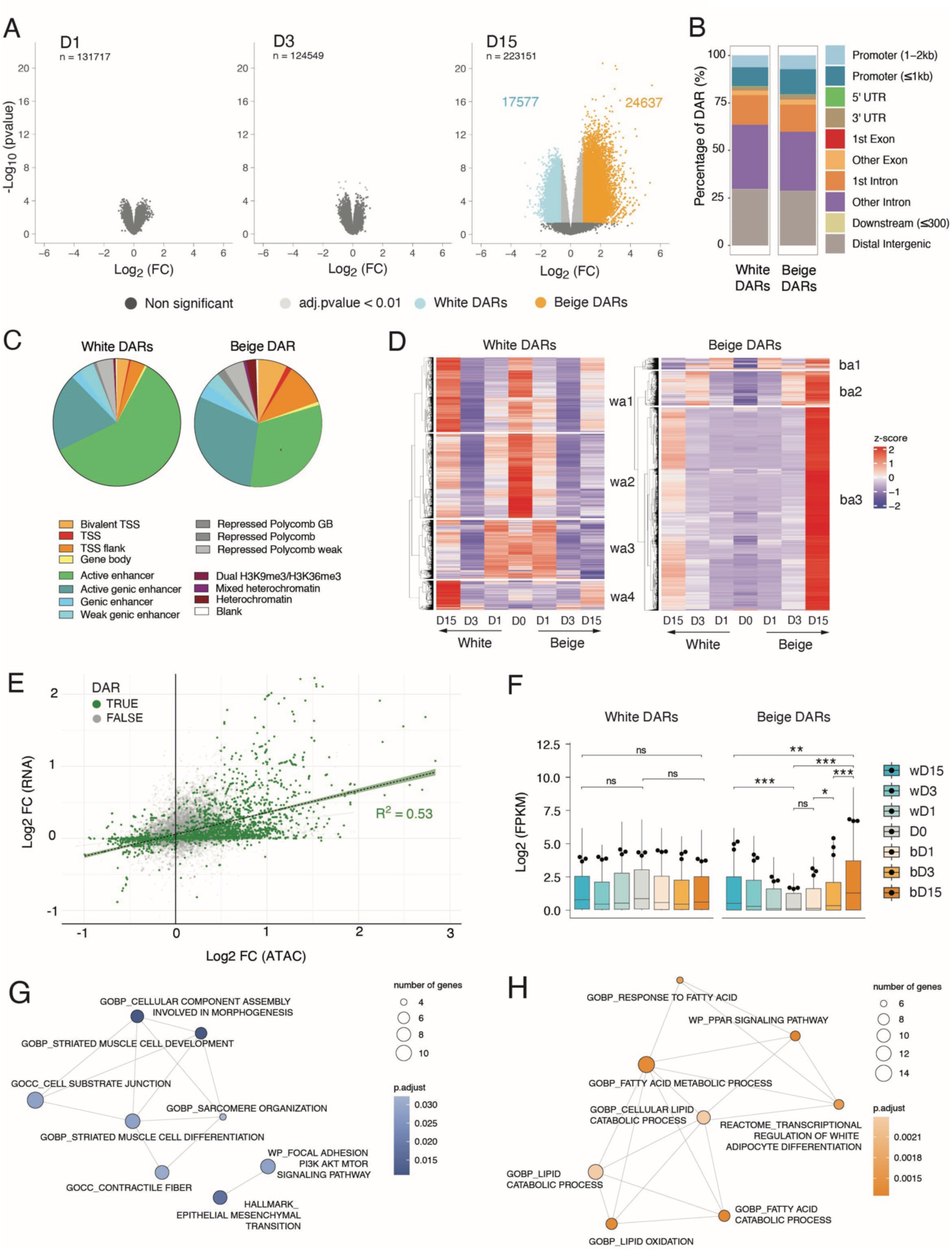
Late opening of promoter regions in beige adipocytes. **A** Volcano plot of differentially accessible regions in white (wDAR) *vs*. beige (bDAR) adipocytes at each time point; n = total number of regions. The number of wDARs and bDARs identified at D15 is indicated on the plot. **B** Annotation of w- and bDARs to genomic regions. **C** Chromatin state annotation of DARs, as percentage of genome coverage. **D** Heatmap with hierarchical clustering of ATAC read counts in DARs, scaled across the differentiation time-course. **E** Scatter plot of log2 fold change ATAC signal versus log2 fold change RNA FPKM at promoters (TSS +/- 2kb) of differentially expressed genes (D15; p < 0.01). Promoters with significant DARs are shown in green. **F** Expression level (log2 FPKM) of genes with differentially accessible promoters across the time-course (white DAR-associated promoters: 211 genes, beige DAR-associated promoters: 222 genes, * p < 0.05, **p < 0.01, ***p < 0.001, t-test with Holm adjustment; ANOVA p < 0.05; n=3 RNA-seq replicates). Boxplots represent the spread of expression between genes. Network of annotated terms enriched for genes with increased accessibility in white (**G**) or beige (**H**) differentiated adipocytes (D15).

We then asked whether D15 DARs arose from ‘closing’ of early-open regions, or from *de novo* ‘opening’ of previously closed or less accessible regions. We assessed the variation in ATAC signals inside D15 DARs across white and beige differentiation (**Fig. 2D**). In agreement with our gene expression data (see **Fig. 1F**), we find that white DARs mainly arise from a reopening of regions that are highly accessible at D0 but transiently close after differentiation induction (**Fig. 2D**, left; clusters wa1-3); only few arise from late opening regions (cluster wa4). In contrast, virtually all beige DARs arise from late (post-D3) chromatin opening events (**Fig. 2D**, right, clusters ba1-3). Quantification of ATAC signals within clusters and at each timepoint corroborates these observations (**Supplemental Fig. S6**).

Since beige DARs are enriched at promoters, we next examined the relationship between chromatin accessibility and gene expression levels. We observe a positive correlation between differential gene expression and differential ATAC-seq signal (R^2^ = 0.53, p < 2.2e^-16^) (**Fig.2E**). In addition, beige DAR-associated genes are strongly induced in bD15 adipocytes, whereas white DAR-associated genes are expressed throughout adipogenesis (**Fig. 2F**). Similar to white DEGs (**Fig.1G**), white DAR-associated genes are implicated in extracellular matrix organization and cytoskeleton interactions (**Fig. 2G)**, while beige DAR-associated genes instead relate to fatty acid metabolic processes (**Fig. 2H**). Altogether, our results indicate that increased accessibility of promoter and enhancers in late differentiation shape the beige adipocyte fate.

### Distinct epigenetic modules contribute to white and beige-specific transcriptional programs

Changes in chromatin accessibility at promoters can be driven by histone modifications and/or differential transcription factor binding and activity. To assess the relative contribution of epigenetic modifications and chromatin opening at promoters to the regulation of gene expression in adipocytes, we established a machine learning model using LightGBM (Ke et al. 2017), which we interpreted with the Shapley additive explanations (SHAP) method (Lundberg and Lee 2017) (**Fig. 3A, Supplemental Fig. S7A-C**). Training on epigenetic features, the model achieves a correlation r^2^ of 0.61 on hidden data **(Supplemental Fig. S7D**) and accurately predicts approximately 70% of variably expressed transcripts ( |log2 FC| > 0.5 and |error| < 0.5; 823 bDEGs and 182 wDEGs) (**Fig. 3B**). However, the model fails to predict changes in TSS expression for a subset of transcripts (1976 transcripts with |log2 FC| > 0.5 and |error| > 0.5), suggesting these are regulated by additional features not included in the model, such as enhancer regulation (**Fig. 3B**; see next section). We thus focused on transcripts for which epigenetic modifications at promoters successfully predicts expression outcome and performed SHAP analysis to explain the contribution of each epigenetic feature to the model prediction. Strikingly, SHAP analysis highlights different modes of regulation for white and beige DEGs promoters, the strongest feature for the prediction of TSS expression being H3K27ac for wDEGs and H3K4me3 for bDEGs (**Fig. 3C**). Indeed, SHAP partial dependence plots highlight distinct relationships between the enrichment of epigenetic features and their influence on model output (**Fig. 3D**): (i) increased H3K4me3 levels are predictive of higher gene expression in beige adipocytes, while this relationship is lost for wDEGs; (ii) the relationship between H3K27ac levels and SHAP values indicate a strong impact for wDEGs and a more moderate effect on model prediction for bDEGs; (iii) the full range of chromatin accessibility signal is linearly correlated with the predictive power of our model, consistent with our previous correlation result (**Fig. 3D**, see **Fig. 2E**). When assessing modes of regulation for each transcript, we confirm that H3K27ac levels are strong predictors for most wDEG expression (**Fig. 3E**, cluster 1 and 3), although changes in chromatin accessibility also contribute to a subset of transcript prediction (**Fig. 3E**, cluster 1). Notably, GO term analysis highlights pathways-specific modes of regulation, with extracellular matrix genes being controlled by both variation in H3K27ac and chromatin accessibility (**Fig. 3F**). For bDEGs, changes in H3K4me3 are the strongest predictor for most transcripts, in association with variations in H3K27ac levels and chromatin accessibility (**Fig. 3C**; **Fig. 3G**, clusters 1,2,4). GO term analysis indicates the increase in mitochondrial genes expression in beige adipocytes is closely tracked by H3K4me3 signal in promoters (**Fig. 3H**). Importantly, the strongest epigenetic predictors according to SHAP analysis also show the most variation at individual TSSs, as exemplified by the *DCN* and *HADH* loci (**Fig. 3I,J**). In summary, our machine learning model points to lineage- and pathway-specific epigenomic regulations of DEG promoters in white and beige adipocytes.

**Figure 3.**
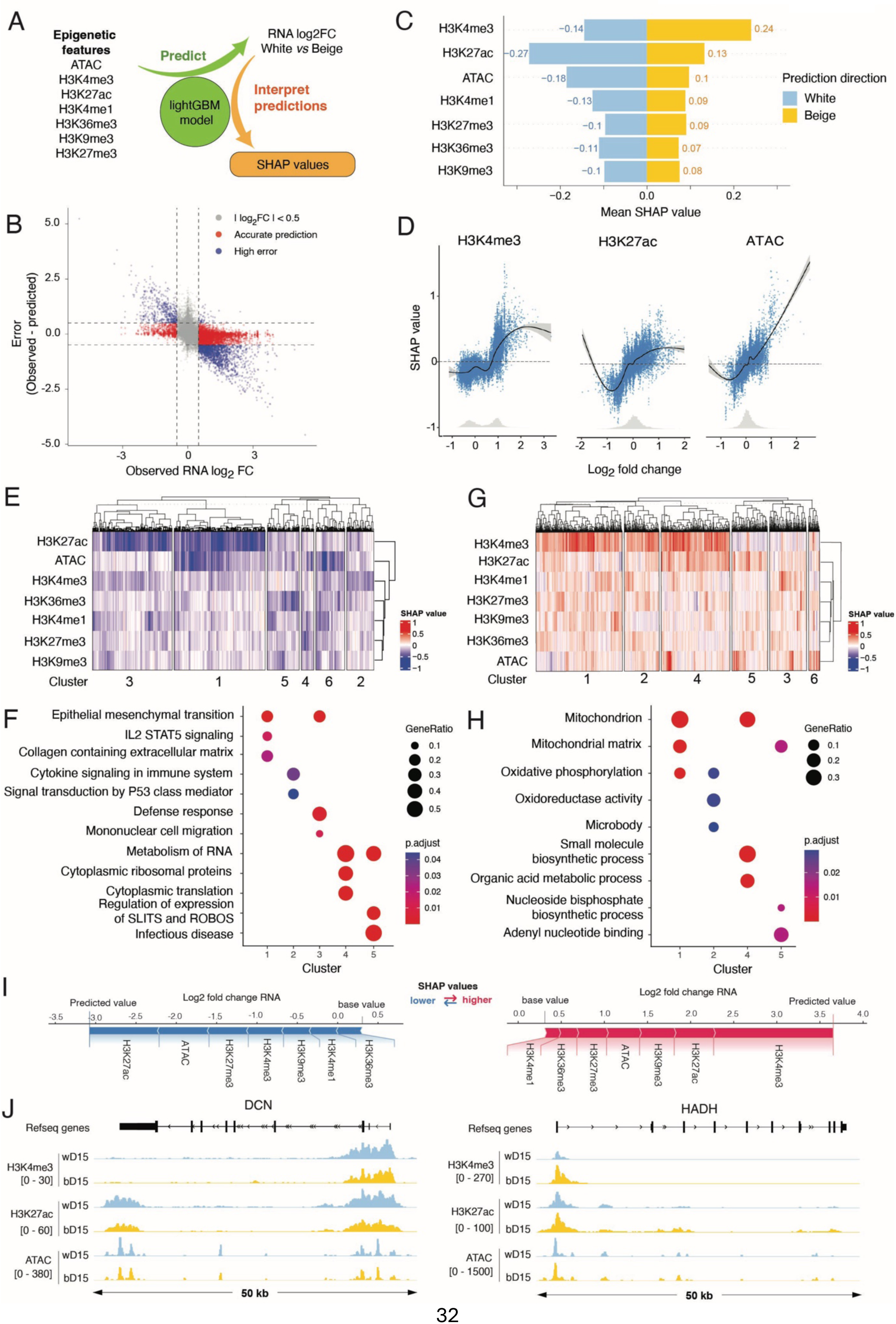
Distinct epigenetic regulations at promoters of white and beige specific transcripts. **A** Methodological flowchart for interpretation of epigenetic datasets using a LightGBM machine learning predictor explained by SHAP. **B** Scatter plot of observed RNA log2 fold change versus prediction error. Transcripts with |RNA log2 fold change | > 0.5 and |error | < 0.5 were considered for further analysis. **C** Average SHAP value for each epigenetic feature stratified by direction of the prediction. **D** SHAP dependency graphs for the predictive power of the H3K4me3, H3K27ac and ATAC features. Grey histograms show the distribution of features values. **E, G** Heatmap with hierarchical clustering of SHAP values for each epigenetic feature for filtered wDEG (**E**) and bDEG (**G**) transcripts. **F, H** Comparative overrepresentation analysis for clusters from **E, G (F,** white SHAP clusters; **H**, beige SHAP clusters) using GO, Reactome and Hallmark gene sets from MSigDb. **I** SHAP waterfall plots showing the effect of epigenetic features on the prediction of the expression of *DCN* (wDEG, left) and *HADH* (bDEG, right). **J** Genome browser views of H3K4me3, H3K27ac and ATAC enrichment at the promoters of white overexpressed (*DCN*: ENST00000052754.10) and beige overexpressed (*HADH*: ENST00000638621.1) genes in white and beige adipocytes.

### Recruitment of short-range enhancers in beige adipocytes

Since changes in promoter epigenetic landscape alone do not explain expression variation for a subset of DEGs transcripts (see **Fig. 3B**), and differential chromatin opening occurs mostly at enhancers (see **Fig. 2C**), we next asked whether dynamic enhancer-promoter interactions also contribute to gene expression changes in white *vs*. beige adipocytes. To capture enhancer interactions at D15, we performed H3K27ac Hi-ChIP (**Supplemental Figure S8A**). When calling chromatin interactions (loops from hereon) using the MAPs pipeline (Juric et al. 2019), we found a similar number of loops across conditions (counts **≥** 6; **Supplemental Figure S8B**), and a complete overlap of the called loops between biological replicates (**Supplemental Figure S8C**). We then called differential loops between white and beige adipocytes using HiC-DC+ (Sahin et al. 2021) (p < 0.05) (**Fig. 4A; Supplemental Table S3**). Amongst 98013 loops called, 3246 loops are significantly enriched in beige adipocytes (”beige loops”) and only 41 loops are enriched in white adipocytes (**Fig. 4A**). Thus, the D15 white-specific transcriptomic profile is not mediated by distinct H3K27ac enhancer interactions.

**Figure 4.**
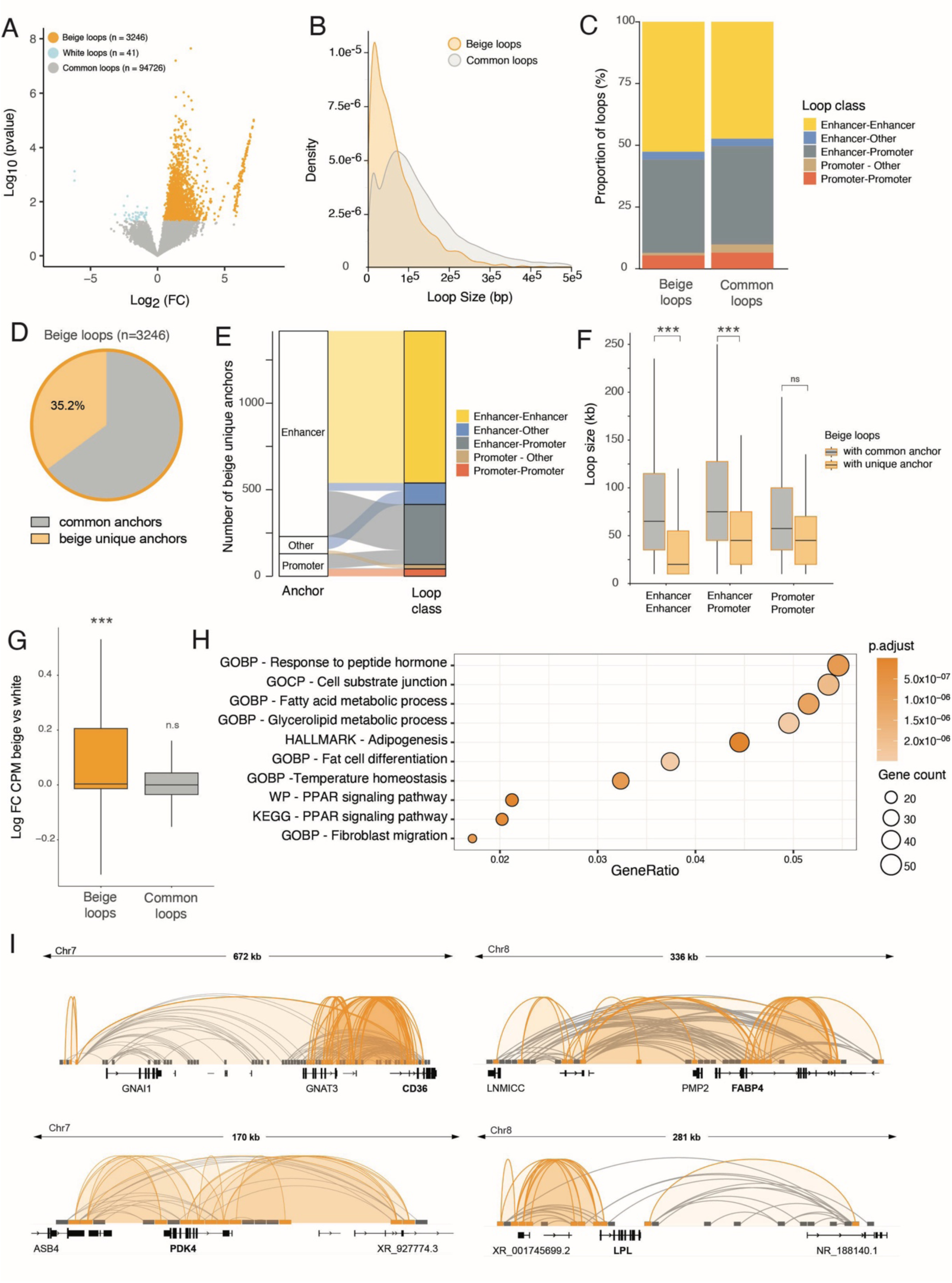
De novo looping of short-range enhancers promote beige gene expression. **A** Volcano plot of differential H3K27ac Hi-ChIP signal in beige *vs.* white adipocytes (D15) (n=3246 beige loops and n=41 white loops, p<0.05). **B** Density distribution of H3K27ac Hi-ChIP loop size, defined as the minimum distance between two anchors. **C** Proportion loop category for H3K27ac Hi-ChIP beige specific and common loops. Anchors located upstream of a TSS (-2kb) and overlapping with a H3K4me3 ChIP peak are defined as Promoter. Anchors overlapping with a H3K27ac ChIP peak are defined as Enhancer. All remaining anchors are defined as Other. **D** Proportion of beige loops with anchors representing *de novo* genomic interactions. **E** Alluvial plot showing the proportion of Enhancer, Promoter and Other categories in *de novo* beige anchors and their corresponding loop annotation. **F** Size of beige loops involving common anchors or beige unique anchors for the indicated loop annotations (*** p < 0.0001, ns non-significant, Wilcoxon rank-sum test) **G** Relative expression (log2 fold change CPM beige *vs.* white, D15) of transcripts associated with beige and common loops. **H** Overrepresentation analysis of genes associated with beige loops using GO, Reactome and Hallmark gene sets from MSigDb. **I** Genome browser views of H3K27ac Hi-ChIP beige (orange) and common (grey) loops at CD36, FABP4, LPL, and PDK4 loci.

Importantly, all beige and common loops overlap with H3K27ac peaks from our D15 ChIP-seq at one or both anchors (**Supplemental Fig. S8D**), validating the Hi-ChIP experiment. Beige loops represent robust interaction events, with more than 40 counts per loop on average (**Supplemental Fig. 8E**). Beige and common loop size ranges from 10 kb to 500 kb, typical for interaction of regulatory elements (Jin et al. 2013) (**Fig. 4B**). However, beige loops show a strong enrichment for shorter range interactions, which could reflect formation of proximal promoter-enhancer loops or the formation of enhancer hubs (**Fig. 4B**). To annotate the loops, we defined 5 classes of interactions based on the presence of H3K4me3 (Promoter) or H3K27ac (Enhancer) peak from our ChIP-seq data within the loop anchors. Enhancer-enhancer and enhancer-promoter interactions are predominant in our dataset, as expected for a H3K27ac Hi-ChIP, with beige loops showing a slight enrichment for enhancer-enhancer interactions (**Fig. 4C**).

We next assessed whether beige loops arise from new interactions between anchors already involved in common loops, or from the recruitment of new genomic regions. Strikingly, 35% of beige loops are formed by *de novo* chromatin contacts (**Fig. 4D**). These beige unique anchors mainly represent enhancer regions, which predominantly connect with other enhancers and with promoters (**Fig. 4E**). Beige-specific enhancer-enhancer and enhancer-promoter interactions formed by *de novo* contacts are significantly smaller than those involving common anchors (**Fig. 4F**). Thus, beige adipogenesis results in both the recruitment of short-range enhancers to promoters, and in the clustering of enhancers in close genomic proximity.

Increased contacts at promoters in beige loops associate with a significant upregulation of expression of the associated genes, in contrast to genes implicated in common loops, which are not differentially expressed in white *vs*. beige adipocytes (**Fig. 4G**). Genes associated with beige loops are enriched for “Fatty acid metabolic processes”, “PPARG signaling pathway” and “Temperature homeostasis” annotations (**Fig. 4H**). Indeed, we detect a strong increase in chromatin contacts at the promoters of genes involved in lipid transport (*LPL*, *CD36*, *FABP4*) and fatty acid oxidation (*PDK4*) (**Fig. 4I**). We conclude that the establishment of the beige adipocyte phenotype is mediated by the recruitment of short range enhancers to regulate the expression of fatty acid metabolism and thermogenic genes.

### Beige loops are enriched in early adipogenic transcription factor binding sites

*De novo* chromatin interactions can be promoted by changes in chromatin state and accessibility, as well as differential binding of TFs and mediator complexes (Dekker and Mirny 2024). We first reasoned that increased chromatin contacts in beige adipocytes detected by H3K27ac Hi-ChIP could be mediated by an increase in H3K27ac levels and/or chromatin opening at loop anchors. We find that nearly half of beige loops harbor a beige DAR at either anchor, and 24.9% display both increased chromatin opening and increased H3K27ac levels (**Fig. 5A**). However, 41.1% of beige loops are formed without significant variations in H3K27ac or ATAC signals at their anchors (**Fig. 5A**), suggesting that differential binding of protein complexes may drive beige-specific 3D genome folding at these sites (Stadhouders et al. 2019).

**Figure 5.**
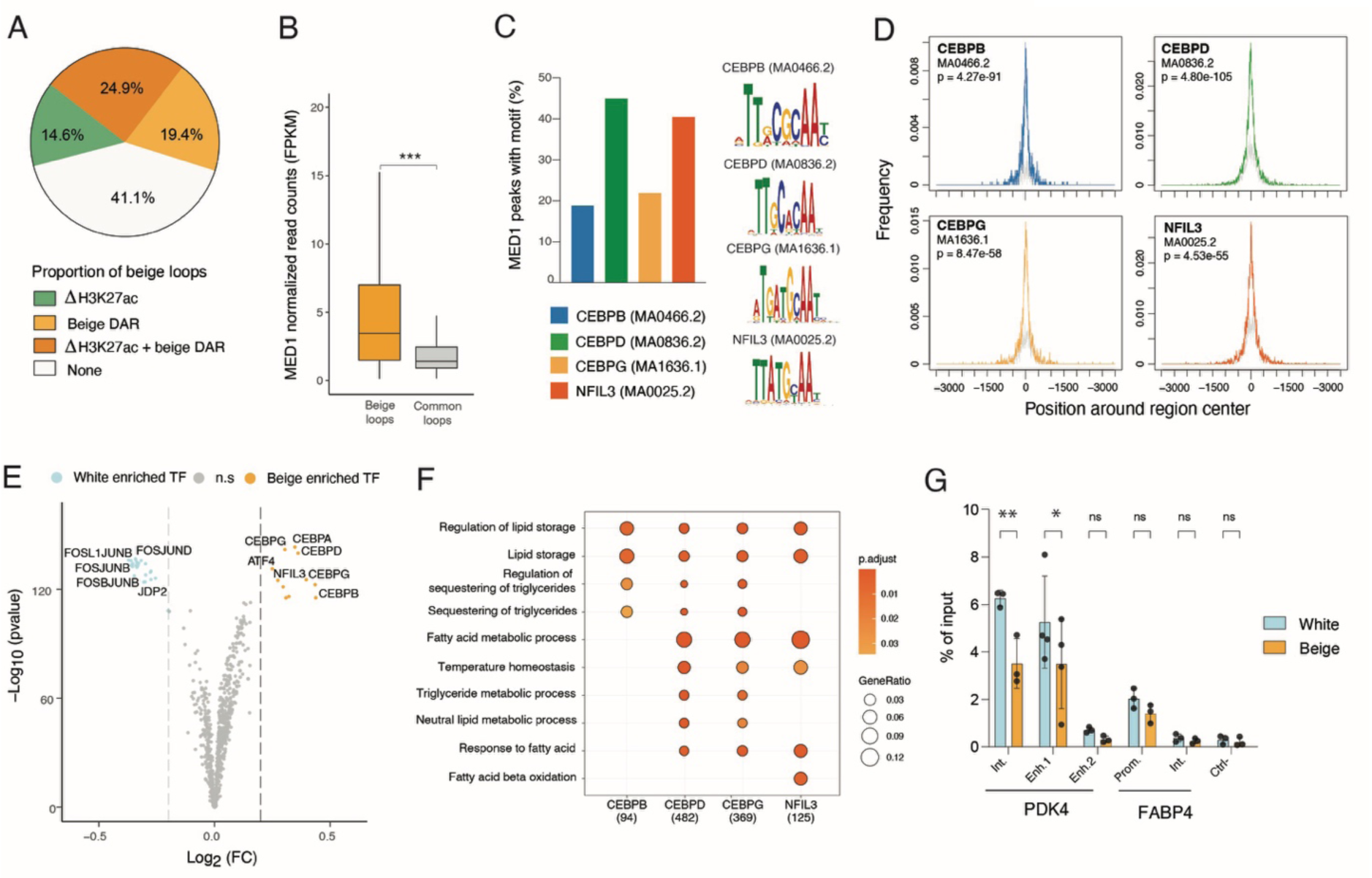
Beige loops are enriched for early adipogenic transcription factor footprints. **A** Proportion of loops with differential H3K27ac peaks (log2 beige *vs.* white fold change > 1) and/or beige DARs at either anchor. **B** Normalized read counts of MED1 ChIP signal (Loft et al. 2015) in beige loops and common loops anchors (*** | D | = 0.98, Cohen’s D standardized mean difference). **C, D** Enrichment analysis for TF motifs within MED1 peaks in beige loops anchors. For the top 4 enriched motifs, proportion of MED1 peaks with motif (**C)** and metaplot of TF motif enrichment relative to genomic regions centers (**D**). **E** Volcano plot of TF footprint enrichment in beige loop anchors in white versus beige D15 adipocytes (white enriched: log2 fold change < - 0.2; beige enriched: log2 fold change > 0.2). **F** Comparative overrepresentation analysis of indicated TF target genes using GO, Reactome and Hallmark gene sets from MSigDb. **G** ChIP-PCR analysis of NFIL3 binding at PDK4 intronic and distal enhancers, FABP4 promoter and intronic region and a negative control region (** p < 0.005, * p < 0.05, ns non-significant, two-way ANOVA with Sidák’s multiple comparison test).

The MED1 subunit has been shown to mediate ligand-dependent binding of the Mediator coactivator complex to nuclear receptors (Chen and Roeder 2011), promoting looping of enhancer communities to promoters in various cellular contexts, including during white and beige adipogenesis (Kagey et al. 2010; Step et al. 2014; Loft et al. 2015; Ito et al. 2021). We thus re-analyzed previously published MED1 ChIP-seq datasets in human beige adipocytes (Loft et al. 2015) and assessed MED1 ChIP signal enrichment at beige and common loops (**Fig. 5B)**. We find significantly higher MED1 levels at beige versus common loop anchors (Cohen’s D = 0.98; **Fig. 5B**), consistent with a higher proportion of active enhancers in beige loops, driving high levels of gene expression (see **Fig. 4C, G**) (Whyte et al. 2013).

To identify TFs potentially associated with MED1 at beige loops, we performed a motif analysis at beige MED1 peaks at beige anchors. We find a significant enrichment for TFs from the C/EBP family, with motifs for C/δ and the C/EBP-related factor NFIL3 being detected in about 40% of the peaks (**Fig. 5C,D**). To predict differential TF binding in white and beige adipocytes, we conducted a differential footprinting analysis on beige loops anchors using ATAC-seq from D15 white and beige adipocytes (**Fig. 5E; Supplemental Table S4**). Strikingly, top enriched TFs at beige loops are members of the C/EBP family, including NFIL3, and their binding partner ATF4 (**Fig. 5E**). In white adipocytes, the same regions are enriched for TFs from the AP-1 family (FOS, FOSL1, FOSL2, JUNB, JUND), and for JDP2, an AP-1 component that potently inhibits AP-1 mediated transcription (Aronheim et al. 1997). Differential footprints for C/EBPs and NFIL3 TFs are found at enhancer-enhancer, enhancer-spromoter and promoter-promoter loop anchors (**Supplemental Fig. 9A**). Genes associated with these differential footprints annotate to fatty acid metabolic processes and temperature homeostasis (**Fig.5F**), and show a global increase in expression in beige *vs.* white adipocytes (**Supplemental Fig. 9B**).

NFIL3 has been described as both a transcriptional repressor and activator (Cowell et al. 1992; Zhang et al. 1995) and can compete with C/EBPs for DNA binding (Li et al. 2011; MacGillavry et al. 2011). Since footprinting analysis cannot discriminate TFs with similar motifs (see **Fig. 5C**), we directly assessed NFIL3 binding profile by ChIP-PCR. NFIL3 is expressed at similar levels between white and beige adipocytes (**Supplemental Fig. 9C, D**). In white adipocytes, we detect a strong binding of NFIL3 at PDK4 regulatory regions, and at the promoter of FABP4 (**Fig.5G, Supplemental Fig.10**). Strikingly, NFIL3 binding is significantly decreased at both intronic and distal enhancer of PDK4 in beige adipocytes (**Fig.5G**). Thus, decreased NFIL3 binding correlates with increased chromatin opening at those sites, increased loop formation, and transcriptional activation. Collectively, these results suggest that increased footprints at *PKD4* enhancers rather reflect an increased binding of C/EBPδ and/or C/EBPβ, and point to NFIL3 as negative regulator of the beige adipocyte phenotype.

## Discussion

Adipocytes are a functionally heterogeneous cell type, and the relative proportion of distinct adipocyte subtypes within adipose tissue associates with differential metabolic disease risk (Reinisch et al. 2024). Beige adipocytes are particularly plastic, as these can convert between the white and beige states in response to environmental changes. Here, we show that the beige adipocyte transcriptomic signature is established late during adipogenesis, and associates with beige-specific chromatin opening at both promoters and enhancers. Distinct epigenetic modulations at promoters drives the upregulation of white- and beige-specific pathways, with beige-specific upregulation of mitochondrial genes correlating with an increase in H3K4me3 levels. In contrast, the upregulation of thermogenic genes cannot be explained by epigenetic remodeling only, but associates with the recruitment of short range enhancers enriched for C/EBP TFs binding motifs.

The establishment of a beige specific thermogenic program associates with a global opening of the chromatin at the promoters of beige-induced genes. We used machine learning and Shapley values to cluster promoters based on the contribution of epigenetic features to changes in TSS expression. This approach allowed us to integrate data across modalities, investigate correlations at each promoter in an error-aware manner, and to overcome potential limitations of feature-specific analysis. We highlight distinct epigenetic modules regulating white and beige transcriptional programs, with respectively higher predictive power for H3K27ac or H3K4me3. Intriguingly, altered levels of these epigenetic marks associates with persistently impaired adipocyte function in a model of weight loss (Hinte et al. 2024). It is thus tempting to speculate that changes in the epigenetic landscape at beige-specific gene promoters could similarly encode a memory of beiging, by either facilitating their reactivation or priming for enhanced transcription upon adrenergic stimulation (Merlin et al. 2018). By measuring the prediction error from our machine learning model, we inferred gaps in the model such as the influence of enhancer looping. While the promoter-enhancer looping feature could in theory be incorporated into the model, the resolution of the Hi-ChIP dataset is ill-suited for the prediction of expression at single TSSs.

Strikingly, our Hi-ChIP analysis of white *vs.* beige enhancer interactions reveals only a negligible proportion of total loops enriched in the white condition, compared to a markedly higher enrichment of beige specific loops. This suggests that the white adipocyte phenotype constitutes the default adipose fate for subcutaneous adipose tissue-derived ASCs, while additional stimuli are needed to acquire the beige phenotype. This is consistent with *in vivo* data showing that beige adipocytes revert to a white phenotype upon stimulus removal (Rosenwald et al. 2013; Altshuler-Keylin et al. 2016). Interestingly our findings suggest that the increase in enhancer looping in beige adipocytes is largely independent of changes in H3K27ac levels, indicating that these regulatory regions are pre-established in white adipocytes but engage in *de novo* interactions through TF recruitment upon beiging stimulation. In particular, both motif and ATAC footprinting analysis point to an enrichment of TFs from the C/EBP family at beige loops anchors. This is consistent with the described role of C/EBPβ as a regulator of beige adipocytes thermogenic program (Kajimura et al. 2009). In contrast, our ChIP-qPCR analysis identifies NFIL3 as a negative regulator of PDK4 expression, a key metabolic enzyme that redirects glucose from oxidation towards triglyceride synthesis, thus favoring the use of fatty acids as energy source (Barquissau et al. 2016; Pettersen et al. 2019). Indeed, we find that decreased NFIL3 binding at PDK4 enhancers correlates with increased transcription in beige adipocytes. Our results notably contradict a recent finding suggesting a positive role for NFIL3 in cAMP-induced adipocyte beiging (So et al. 2022). NFIL3 can act as both a transcriptional activator and repressor (Cowell et al. 1992; Zhang et al. 1995). For instance, NFIL3 can heterodimerize with other bZIP TFs such as CREB to form context-specific transcriptional complexes (Acharya et al. 2006). However, it can also repress genes activated by C/EBPs by competing for the same motif (MacGillavry et al. 2011; Chen et al. 2024). Thus, the apparent discrepancy in the role of NFIL3 between rosiglitazone- and cAMP-induced beiging systems might stem from the context-specific actions of NFIL3.

It is important to acknowledge that our findings are largely correlative, and future functional experiments are needed to establish a causal relationship between genome remodeling and the establishment of a beige phenotype. In addition, given the timescale of adipose differentiation, this experimental setup lacks the temporal resolution needed to infer causality direction between (epi)genome remodeling and transcriptional changes (Reed et al. 2022). Indeed, we cannot rule out that transcriptional activation is the cause, rather than the result, of differential loop formation. Nevertheless, our results highlight the complex epigenetic and enhancer network remodeling associated with the establishment of the beige adipocyte identity.

## Materials and Methods

### Cell culture and adipose differentiation

Primary ASCs were isolated from subcutaneous fat obtained by liposuction from a female donor (age 45; BMI 20,9 kg/m^2^), obtained after donor’s informed consent as approved by the Regional Committee for Research Ethics for Southern Norway with number REK 2013/2102. ASCs were cultured in DMEM/F12 (17.5 mM glucose) with 10% fetal calf serum and 20 ng/ml basic fibroblast growth factor (proliferation medium). Upon confluency, fibroblast growth factor was removed, and cells cultured for 72 h in DMEM/F12 (17.5 mM glucose) with 10% fetal calf serum (basal medium) before induction of differentiation (day 0: D0). For white adipose differentiation, ASCs were induced with a cocktail of 0.5 µM 1-methyl-3 isobutyl xanthine (Sigma, I5879) , 1 µM dexamethasone (Sigma, D4902), 10 µg/ ml insulin (Sigma, I9278) and 200 µM indomethacin (Sigma, I7378-56) in basal medium. Differentiation media was renewed every 3 days until day 9, after which cells were maintained in DMEM/F12 (17.5 mM glucose) with 10% fetal calf serum and 10 µg/ ml insulin. For beige adipose differentiation, media were supplemented with 1 µM Rosiglitazone (Sigma, R2408) until day 15. Samples were harvested on D0, then D1, D3 and D15 after induction. All differentiation experiments were done in at least three biological replicates with ASCs at passage 3-8.

### Oil red O staining and quantification

Non-induced ASCs and D15 cells were fixed in 4% paraformaldehyde for 20 min and intracellular neutral lipids were labeled for 30 min with Oil Red-O (O0625; Sigma-Aldrich) diluted in isopropanol. Staining was next eluted in 100% ethanol and absorbance measured at 500nm.

### Microscopy and image analysis

Cells were differentiated for 15 days on 12-mm diameter coverslips in 24-well plates. Cells were washed 3 times with PBS before fixation in 4% paraformaldehyde for 10 min. Cells were then incubated in Bodipy (1µg/ml, Invitrogen D3922) for 15 min and washed 3 times in PBS before mounting in DAKO Fluorescence Mounting Medium (S3023, Agilent) containing 0.2 µg/ml DAPI. Images were acquired on an IX81 microscope (Olympus) fitted with epifluorescence, a 100× 1.4 NA objective mounted on a piezo drive, and a DeltaVision personalDV (Applied Precision, Ltd.) imaging station. For lipid droplet measurements, images were segmented with ImageJ’s Morphological Segmentation plugin MorphoLibJ (https://imagej.net/plugins/morpholibj), and area was measured with the “Analyze particles” function, thresholding on circularity and size.

### Immunoblotting

Proteins were resolved by gradient 4-20% SDS–PAGE, transferred onto nitrocellulose (Cat. 162-0115, BioRad) or PVDF (Cat. IPFL00010, Millipore) membranes and blocked with 5% BSA or non-fat dry milk. Membranes were incubated for 1h at room temperature or overnight at 4°C using the following antibodies: FAS (Santa Cruz, sc-48357), PPARG (Thermofisher, MA5-14889), CD36 (Santa Cruz, sc-9154), CITED1 (Novus, H00004435-M03), UCP1 (Abcam, 23841), NFIL3 (Abcam, EPR27211-70), γTubulin (Sigma, T5326), Total OXPHOS human antibody cocktail (Abcam, ab11041), and GAPDH (Santa Cruz, sc-25778). Proteins were visualized using IRDye-800- (LI-COR Biosciences, 926-32214), IRDye-680- (LI-COR Biosciences, 926-68023), or HRP-coupled (Jackson ImmunoResearch, 111-035-144 and 115- 035-146) secondary antibodies. Bands were quantified by densitometry (BioRad Image Lab) using γTubulin or GAPDH for normalization. Uncropped membranes are presented as **Supplemental Fig. S1 and S3.**

### RNA-seq

Total RNA was isolated from triplicate experiments using the RNAeasy mini kit (Qiagen). Sequencing libraries were prepared using the KAPA mRNA HyperPrep kit (Roche) and sequenced on a Novaseq (Illumina). RNA-seq reads were filtered to remove low-quality reads using fastp v 0.20.1 (Chen et al. 2018). Filtered reads were aligned to the hg38 genome (GENCODE v32) with hisat2 v2.1.0 (Kim et al. 2019), and counted using featureCounts in Subread v2.0.1 with options -M and --fraction (Liao et al. 2014). Low abundance genes were filtered using filterByExpr and then normalized using the trimmed mean of M values (TMM) method from edgeR (Robinson and Oshlack 2010). Beige and white gene expression was compared between conditions using the robust eBayes method with limma-voom adjustment (Law et al. 2014). DEGs had FDR adjusted p-value < 0.01 and absolute log2FC > 1. Heatmaps were generated by hierarchical clustering of expression z-scores using in the R package ComplexHeatmap. To calculate FPKM at promoters, RNA reads were counted 300bp downstream of each TSS using featureCounts with -O and normalization factors were calculated with the TMM method.

### ChIP-seq of modified histones

ChIP of H3K27ac, H3K4me3, H3K4me1, H3K27ac, H3K27me3, H3K36me3, and H3K9me3 was done as described (Rønningen et al. 2015). Undifferentiated cells were fixed for 10 min with 1% formaldehyde, lysed in 50 mM Tris-HCl, pH 8, 10 mM EDTA, 1% SDS, protease inhibitors and Na-sodium butyrate, and sonicated in a Bioruptor Pico (Diagenode) into ∼200 bp fragments. After sedimentation, the supernatant was diluted 10 times in RIPA buffer and incubated with H3K4me3 (Diagenode c15410003), H3K4me1 (Diagenode c15410037), H3K27ac (Diagenode c15410174), H3K27me3 (Diagenode c15410069), H3K36me3 (Diagenode c15410058) and H3K9me3 (Diagenode c15410056) antibodies (each at 2.5 µg/10^6^ cells) coupled to magnetic Dynabeads Protein A (Invitrogen) for 2 h at 4°C. ChIP samples were washed, cross-links were reversed and DNA was eluted for 2 h at 68°C in 50 mM NaCl, 20 mM Tris-HCl pH 7.5, 5 mM EDTA, 1% SDS and 50 ng/µl Proteinase K. DNA was purified using phenol:chloroform:isoamyl alcohol and dissolved in H_2_O. For D15 differentiated white and beige adipocytes, cells were trypsinized, resuspended in HBSS, 0.5% BSA, and centrifuged 200g for 5 min to isolate floating mature adipocytes. Purified nuclei were then fixed and processed as cell samples. ChIP-seq libraries were prepared using a Microplex kit (Diagenode) and sequenced with 150-bp paired-end reads on an Illumina NovaSeq.

### ChIP-seq analysis

FASTQ sequences from H3K4me3, H3K27ac, H3K4me1, H3K36me3, H3K27me3 and H3K9me3 ChIPs were aligned to hg38 using Bowtie2 v2.4.5 (Langmead and Salzberg 2012). Peaks were detected using MACS2 v2.2.7.1 (Zhang et al. 2008), except for H3K9me3 ChIP where peaks were called using Enriched Domain Detector (http://github.com/CollasLab/edd) (Lund et al. 2014) with gap penalty and bin size defined as the mean outputs of 10 runs in auto-estimation mode. ChIP-seq read counts were normalized to library size using reads per genome coverage (RPGC) method in deepTools. Log2(ChIP/Input) ratios were calculated using bamCompare in deepTools v3.5.3 (Ramírez et al. 2014). Bigwig files of normalized read counts were visualized using Integrative Genomics Viewer (Robinson et al. 2011). Differential peaks were identified using MAnorm2 v1.2.2 with default parameters (Tu et al. 2021).

A 15-state chromatin model was generated using chromHMM (Ernst and Kellis 2017) with aligned reads in 1 kb bins. Six histone modifications each from day0, and day15 white and beige samples were included.

### ATAC-seq

ATAC-seq was done following the Omni-ATAC protocol (Corces et al. 2017) and optimized for D15 adipocytes. Cells on D0, D1 and D3 were trypsinized and 50000 viable cells per condition were pelleted at 500 g for 5 min at 4⁰C. Nuclei were isolated by lysing cell pellets in 50 µl of ice-cold RSB buffer (10 mM Tris-HCl, pH 7.4, 10 mM NaCl, 3 mM MgCl_2_, 0.1% NP40, 0.1% Tween-20, 0.01% digitonin) and ending lysis with 1 ml RSB buffer lacking NP40 and digitonin, followed by centrifugation at 500 g for 10 min at 4⁰C. Supernatants were removed and nuclei pellets were carefully resuspended in 50 µl of transposition mixture (25 µl 2X TD buffer and 2.5 µl transposase (Illumina, 20034197), 16.5 µl PBS, 0.5 µl 1% digitonin, 0.5 µl 10% Tween-20, 5 µl H_2_O) to carry out the transposition reaction at 37⁰C for 30 min on a Thermomixer at 1000 rpm. Cells on D15 were washed with PBS and treated 30 min with 200U/ml DNAse I (Worthington Biochemical, LS002139) in DMEM/F12 10% FBS at 37°C. After 4 washes with warm PBS, cells were lysed directly on the culture dish on ice using RSB containing 1% NP40, 0.1% Tween-20 and 0.01% digitonin. After a 5 min centrifugation at 500 g 4°C, pelleted nuclei were resuspended in RSB lacking NP40 and digitonin, then stained with Trypan Blue and counted on a Neubauer chamber. 50000 nuclei were transferred to 1ml of the same buffer to be pelleted during a 10 min centrifugation at 500g 4°C. Tagmentation was performed as above. Total tagmented DNA was purified using the Zymo DNA Clean and Concentrator-5 kit (Zymo Research, D4014) and eluted in 20 µl elution buffer. Libraries were amplified initially for 5 cycles using dual indexes for tagmented libraries from Diagenode (C01011034) with NEB Next High-Fidelity 2x PCR Master Mix (NEB, M0541). Further amplification was adjusted to the input material as assessed by qPCR using primer pair UDI 1 (Diagenode) for all samples. Library clean-up was done twice with a double-sided selection using AMPure XP beads (Beckman Coulter). Cleaned-up libraries were quantified with the KAPA Library Quantification Kit (Roche, KK4824) and quality assessed using the Agilent TapeStation System, before sequencing at > 200 million read depth with 150 bp paired-end reads on an Illumina NovaSeq.

### ATAC-seq analysis and TF footprinting

ATAC-seq reads were filtered by removing low-quality reads using fastp v0.23.2 (Chen et al. 2018). Reads were aligned hg38 with Bowtie2 v2.4.5 (Langmead and Salzberg 2012). Duplicates were removed using Picard MarkDuplicates (http://broadinstitute.github.io/picard/) along with mitochondrial reads and reads that aligned with a mapping quality below 10 using samtools v1.9 (Li et al. 2009). Genrich v0.6.1 was used to call peaks for each sample (https://github.com/jsh58/Genrich). Normalized read count tracks were generated using deepTools. ATAC differential regions were identified using MAnorm2 v1.2.2 with default parameters (Tu et al. 2021). TF footprinting was done using TOBIAS v0.16.0 (Bentsen et al. 2020) with default parameters. JASPAR2022 (CORE_vertebrates_non-redundant) was used for motif enrichment analysis to identify TF binding sites (Castro-Mondragon et al. 2022). DARs were considered promoter-associated when located within 2kb upstream of a TSS harboring a H3K4me3 peak at any differentiation time point.

### Machine learning model and SHAP analysis

The change in seven epigenetic signal tracks (ATAC-seq and ChIP-seq of H3K4me3, H3K4me1, H3K27ac, H3K27me3, H3K36me3 and H3K9me3) between white and beige were used to predict the average RNA log2 fold change at promoters with a lightGBM model. Epigenetic signals were summarized as a log2 fold change white *vs.* beige at promoters of DEGs (TSS -2 kb / +300bp) from RPGC normalized read counts. FPKM at promoters was calculated as indicated above.

Python package lightGBM was used to train a gradient boosting regression tree, predicting the expression of 18 282 promoters from seven epigenetic tracks. To determine the optimal hyper parameters, several values of number of estimators, learning rate and the maximum number of leaves per tree were tested with 10-fold cross validation (**Supplemental Fig. S7)**. With fixed parameters for max_bin and min_data_per_leaf (512 and 100 respectively), model accuracy (measured via negative root mean squared error and r^2^) increased with the number of estimators until about 10, 000 estimators (**Supplemental Fig. S7A, B**). Accuracy also correlated positively with learning rate. Increasing the maximum number of leaves per estimator increased both model accuracy and computational time (**Supplemental Fig. S7B, C**) The final parameters to the model were: number of estimators = 10 000, learning rate = 0.05 and the maximum number of leaves per tree = 50 with 10-fold cross validation. A random forest regression model was fitted to the same data (n_estimators = 10, 000, max_leaf_nodes = 50, min_samples_leaf = 100), which took significantly longer to run and produced significantly worse predictions (**Supplemental Fig. S7E, F**).

### Hi-ChIP and chromatin loop calling

ASCs were differentiated in 10 cm Petri dishes. Cells were washed twice in warm PBS before lysis in 4 ml Hi-ChIP buffer (10mM Tris-HCl pH 7.4, 10mM NaCl, 3 mM MgCl_2_; 0.01% Digitonin, 0.1% Tween 20 and 1.0% NP-40, 20 mM Na-Butyrate). Lysed cells were centrifugated at 500 g for 5 min at 4⁰C. The floating fraction (containing unlysed adipocytes) and the pellet were combined in 5 ml Hi-ChIP buffer and dounced to isolate nuclei. Nuclei were washed twice in Hi-ChIP buffer, followed by centrifugation at 500 g for 5 min at 4⁰C. The nuclei pellet was resuspended in 5 ml Nuclei buffer (10mM HEPES, 1.5 mM MgCl_2_, 250 mM sucrose, 0.1 % NP-40, 20 mM Na-Butyrate in PBS) and processed for Hi-ChIP according Arima-HiC+ Kit protocol (Cat Nr: A510008), with some adaptations. Briefly, 15 µg DNA equivalent of crosslinked nuclei were lysed in deionized H2O, sonicated for 10 cycles (30s on, 30s off; Bioruptor Pico). The ChIP was performed with 2.5 µg H3K27ac antibody (Diagenode C15410174). Libraries were quantified (KAPA, KK4824), amplified (KAPA, KK2620) and barcoded with xGen 2S Plus DNA library prep kit (IDT, 10009877) in combination with xGen 2S MID Adapter (IDT, 10009900). Libraries were sequenced to an average depth of 355 million 150 bp PE reads on an Illumina NovaSeq instrument.

H3K27ac Hi-ChIP was analyzed using MAPS v2.0 (Juric et al. 2019) as implemented within the Arima Genomics pipeline (https://github.com/HuMingLab/MAPS/tree/master/Arima_Genomics). Briefly, the raw paired-end reads were mapped to the hg38 reference genome using bwa mem v0.7.12 (Li et al. 2009), filtered for uniquely mapped reads, then binned at 5 kb to generate the chromatin contact matrix. MACS2 2.2.9.1 was used for H3K27ac peaks calling (Zhang et al. 2008). Paired-end reads with at least one end overlapping the H3K27ac ChIP–seq peaks were used to identify long-range chromatin interactions at 5 kb resolution. A positive Poisson model was used for identifying significant interactions (FDR < 0.01).

HiC-CDC+ (Sahin et al. 2021) was used to call differential loops with the following adaption: significant loops with more than 6 read counts in both replicates and a total count over 18 in at least one of two conditions (white and beige) were included.

### Motif enrichment analysis

Motif enrichment analysis was performed on MED1 ChIP peaks (Loft et al. 2015) overlapping with beige loops anchors against the complete JASPAR database using TFmotifView web server (https://bardet.u-strasbg.fr/tfmotifview/).

### NFIL3 ChIP-PCR

ASCs were differentiated in 10 cm Petri dishes until D15. Nuclei were isolated from approximately 10x10^6^ cells and fixed for 10 min with 1% formaldehyde in Nuclei buffer (10mM HEPES, 1.5 mM MgCl_2_, 250 mM sucrose, 0.1 % NP-40). Fixed nuclei were lysed in 50 mM Tris-HCl, pH 8, 10 mM EDTA, 1% SDS and protease inhibitors, and sonicated in a Bioruptor Pico (Diagenode) into ∼200 bp fragments. After sedimentation, the supernatant was diluted 10 times in RIPA buffer and incubated overnight at 4°C with 8 µg NFIL3 ChIP-grade antibody (Abcam, EPR27211-70), followed by incubation with magnetic Dynabeads Protein A (Invitrogen) for 2h at 4°C. ChIP samples were washed, cross-links were reversed and DNA was eluted for 2 h at 68°C in 50 mM NaCl, 20 mM Tris-HCl pH 7.5, 5 mM EDTA, 1% SDS and 133 ng/µl Proteinase K. DNA was purified using phenol:chloroform:isoamyl alcohol and dissolved in H_2_O. ChIP DNA was used as template for quantitative (q)PCR using primers targeting NFIL3 binding sites and a control region (**Supplemental Table S5**). PCR was done using SYBR Green (BioRad) for 3 min at 95°C and 40 cycles of 95°C for 30 s, 60°C for 30 s, and 72°C for 20 s.

## Competing interest statement

None declared.

## Acknowledgments

We acknowledge the Norwegian Sequencing Centre (Oslo University Hospital) for professional sequencing services.

## Author contributions

SHP and MA analyzed data. NMG and ALS generated datasets. MZ advised on machine learning and statistics. NB and SHP made figures. NB and PC designed the study. NB supervised the work. NB, SHP and PC wrote the manuscript. All authors approved the final version of the paper.

## Funding

This work was funded by the University of Oslo, South-East Health Norway grant 40202 to PC, Research Council of Norway grant 313508 to PC, and Nansen fund 17368 to NB.

## Notes

### Competing Interest Statement

The authors have declared no competing interest.

