## Supplementary information for "Multimodal epigenetic and enhancer network remodeling shape the transcriptional landscape of beige adipocytes"

**Supplemental tables** (Excel)

**Table S1.** Related to Figure 1. Differentially expressed genes in white vs beige adipocytes (D15).

**Table S2**. Related to Figure 2. ATAC peaks summary statistics.

**Table S3.** Related to Figure 4. Hi-ChIP differential loop analysis (HiC-DC+)

**Table S4**. Related to Figure 5. Differential ATAC footprinting analysis (TOBIAS)

**Table S5**. Related to Figure 5. ChIP-PCR primers sequences

**Supplemental figures**

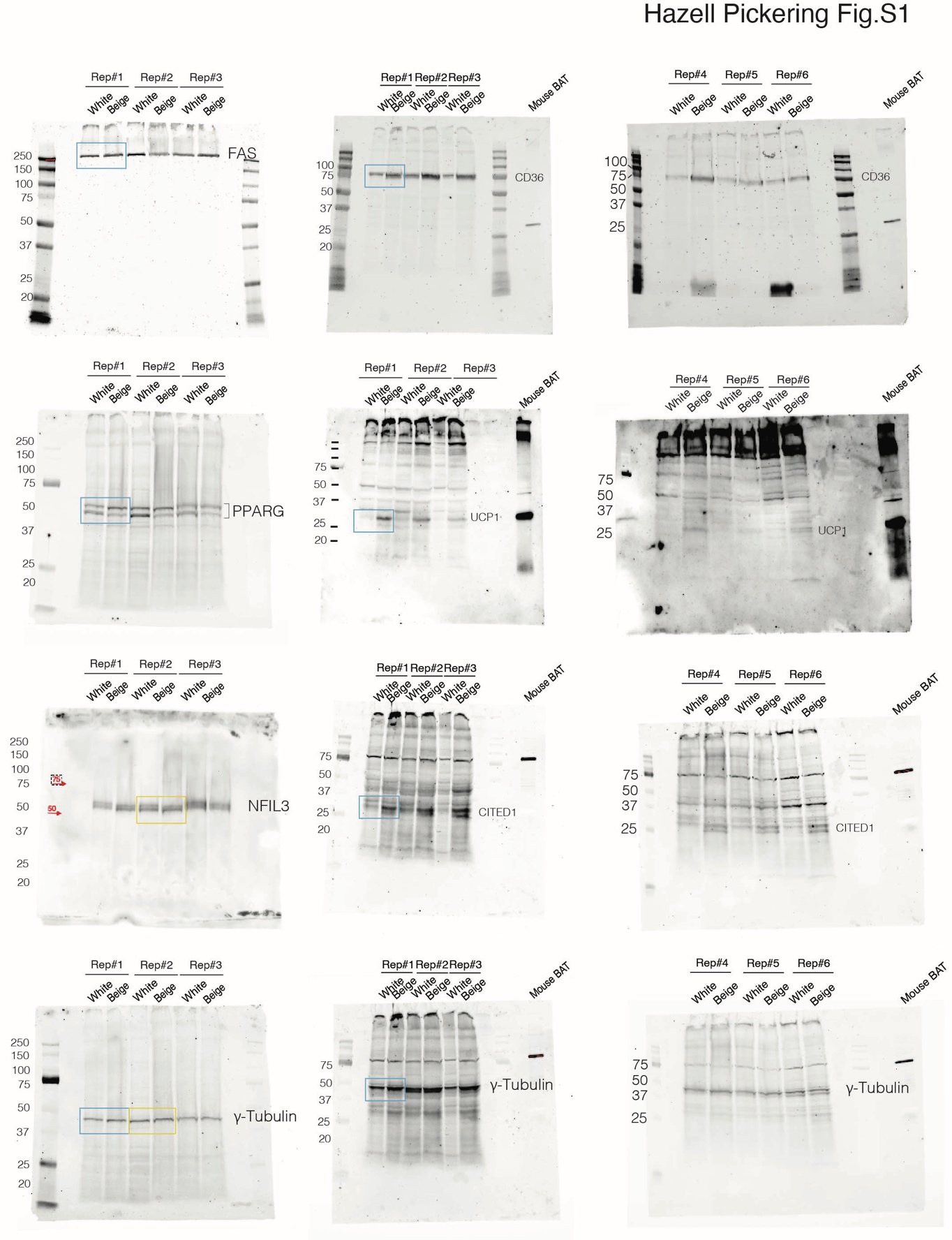

**Fig. S1.** Related to Fig1. Uncropped Western blot replicates for Fig.1 C and S9D.

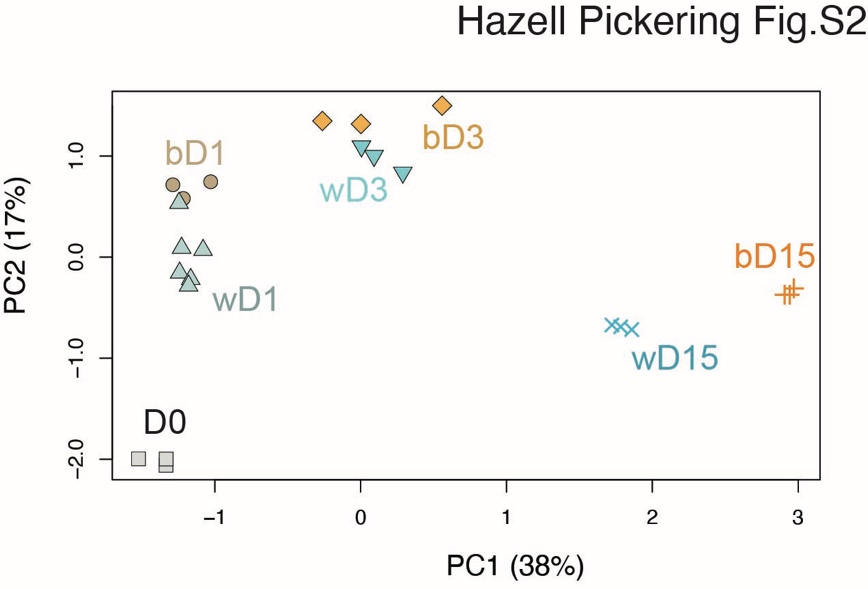

**Fig. S2**. Related to Fig1. Principal component analysis of time course RNA-seq.

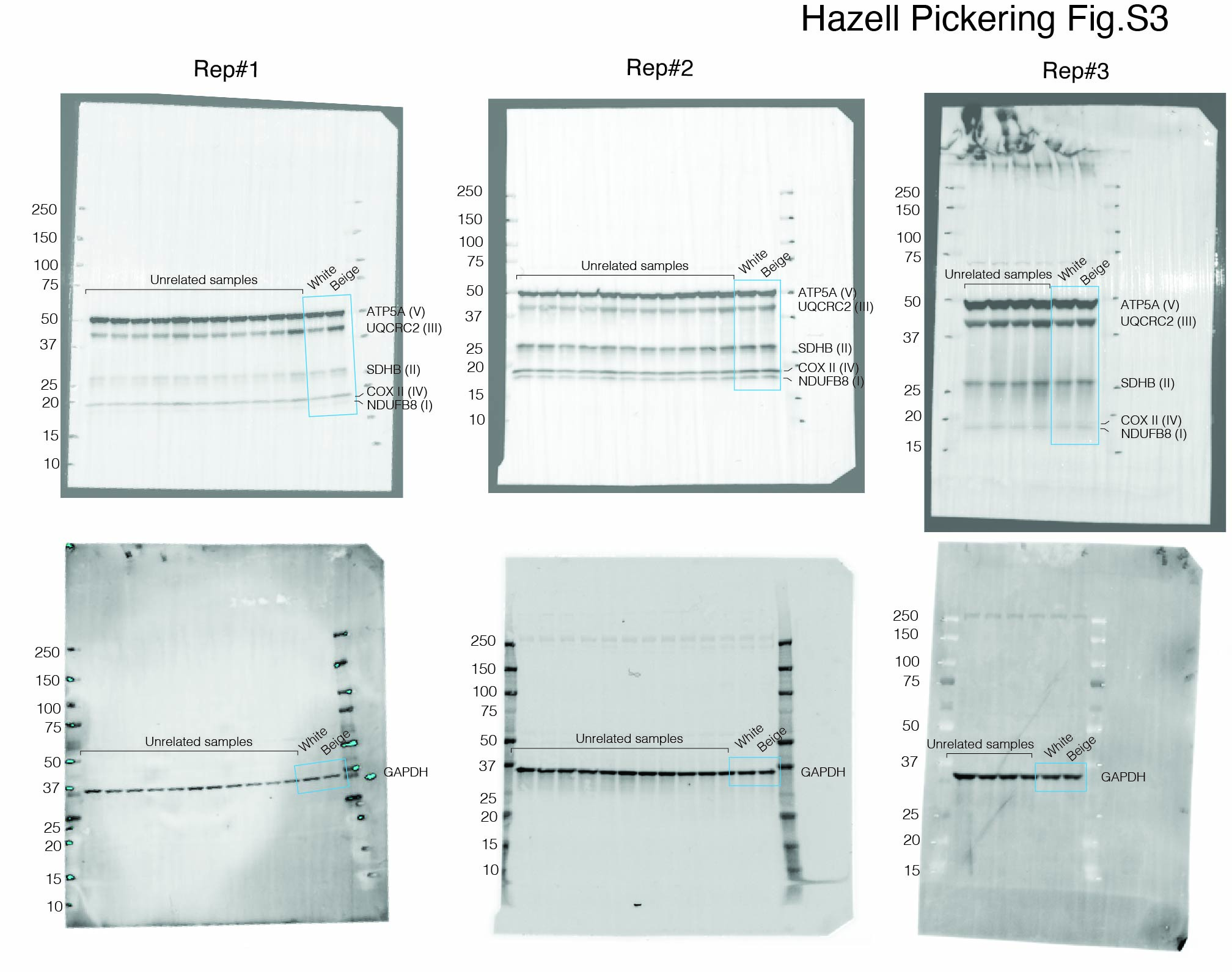

**Fig. S3.** Related to Fig1. Uncropped Western blot replicates for Fig.1 H.

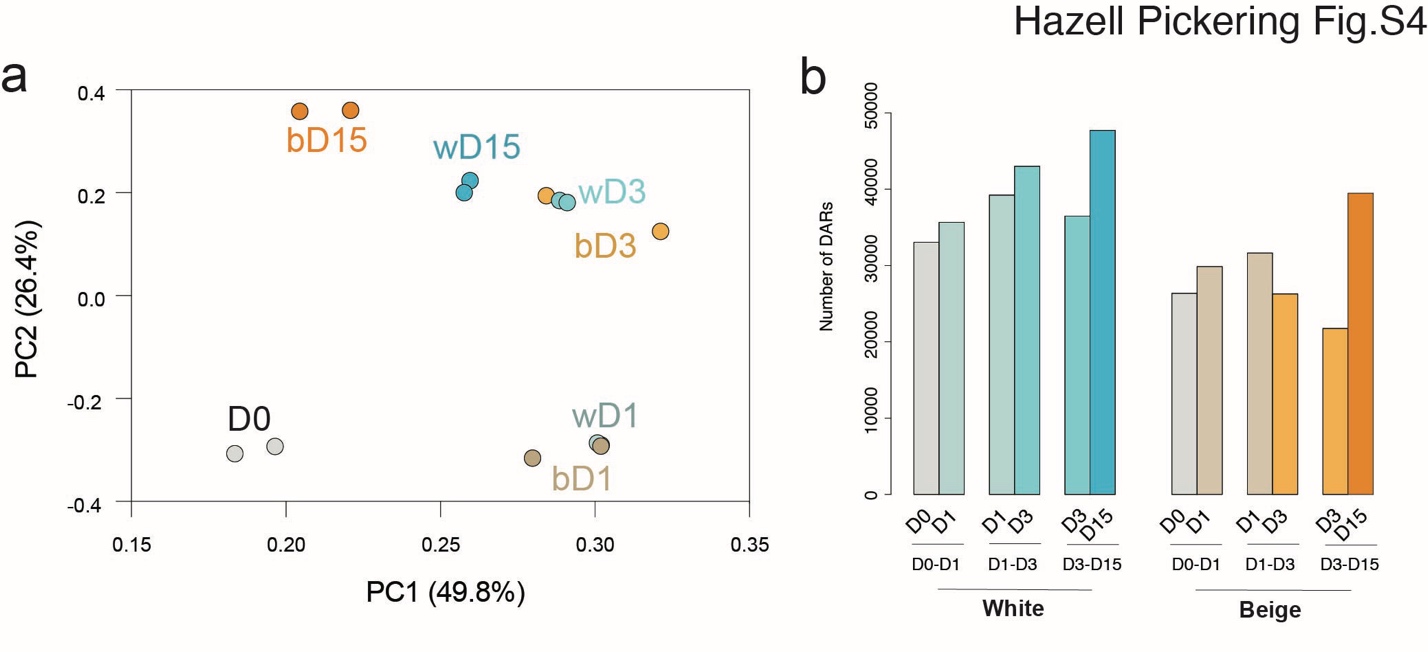

**Fig. S4**. Related to Fig2. **A** principal component analysis of time course ATAC-seq. **B** Number of DARs across the time course.

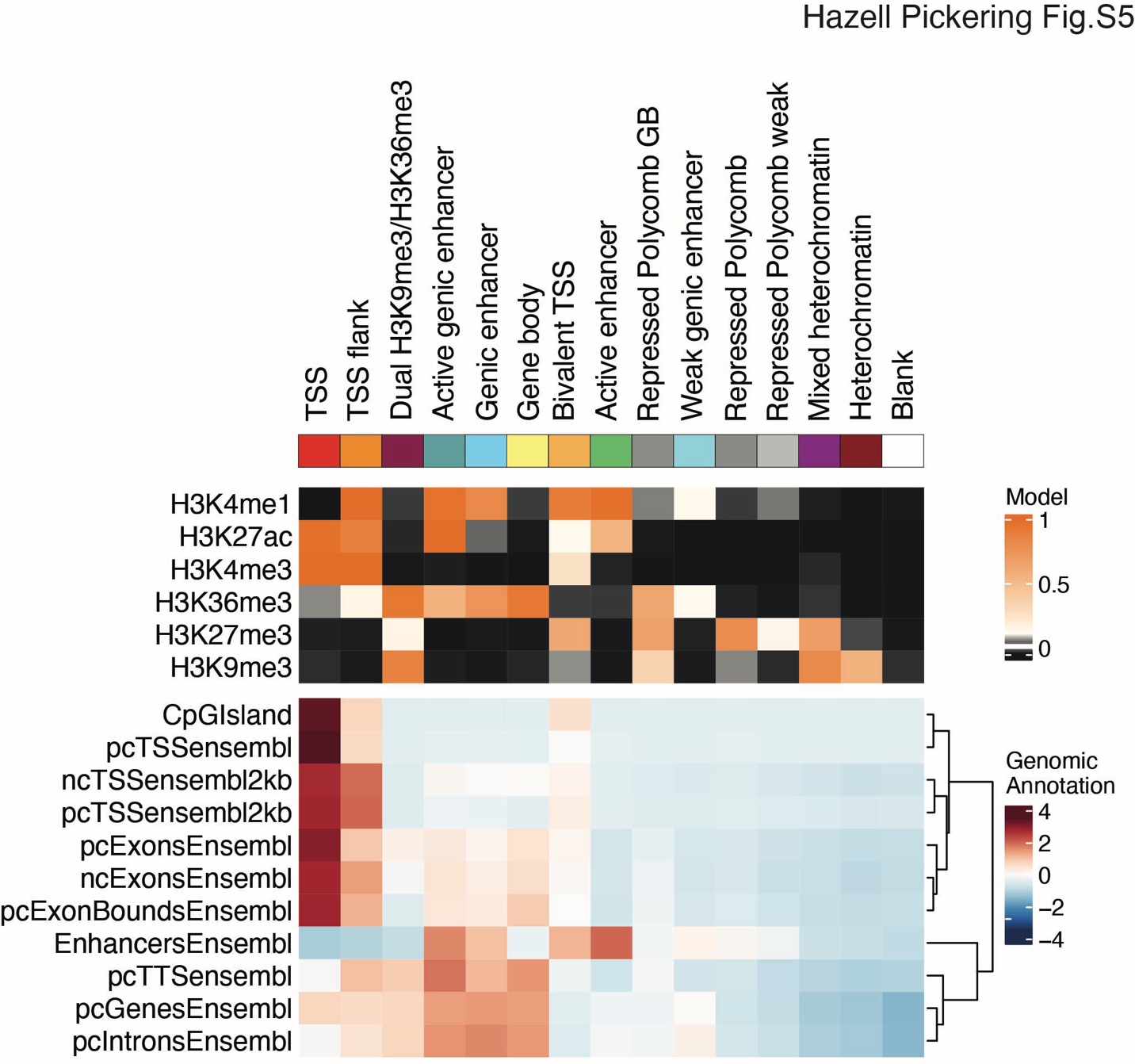

**Fig. S5**. Related to Fig2. Emissions from the 15-state chromatin state model (top panel) and corresponding genomic annotation (lower panel)

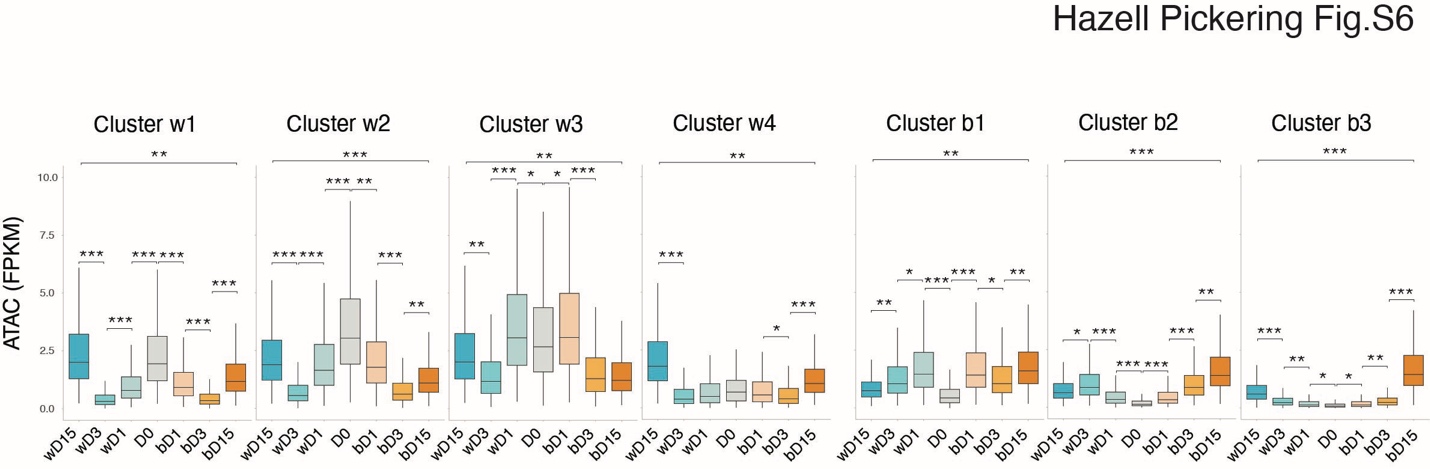

**Fig. S6.** Related to Fig2. Quantification of ATAC signals within clusters (***| D | > 0.8, **| D | > 0.5, *| D | > 0.2; Cohen's D standardized mean difference).

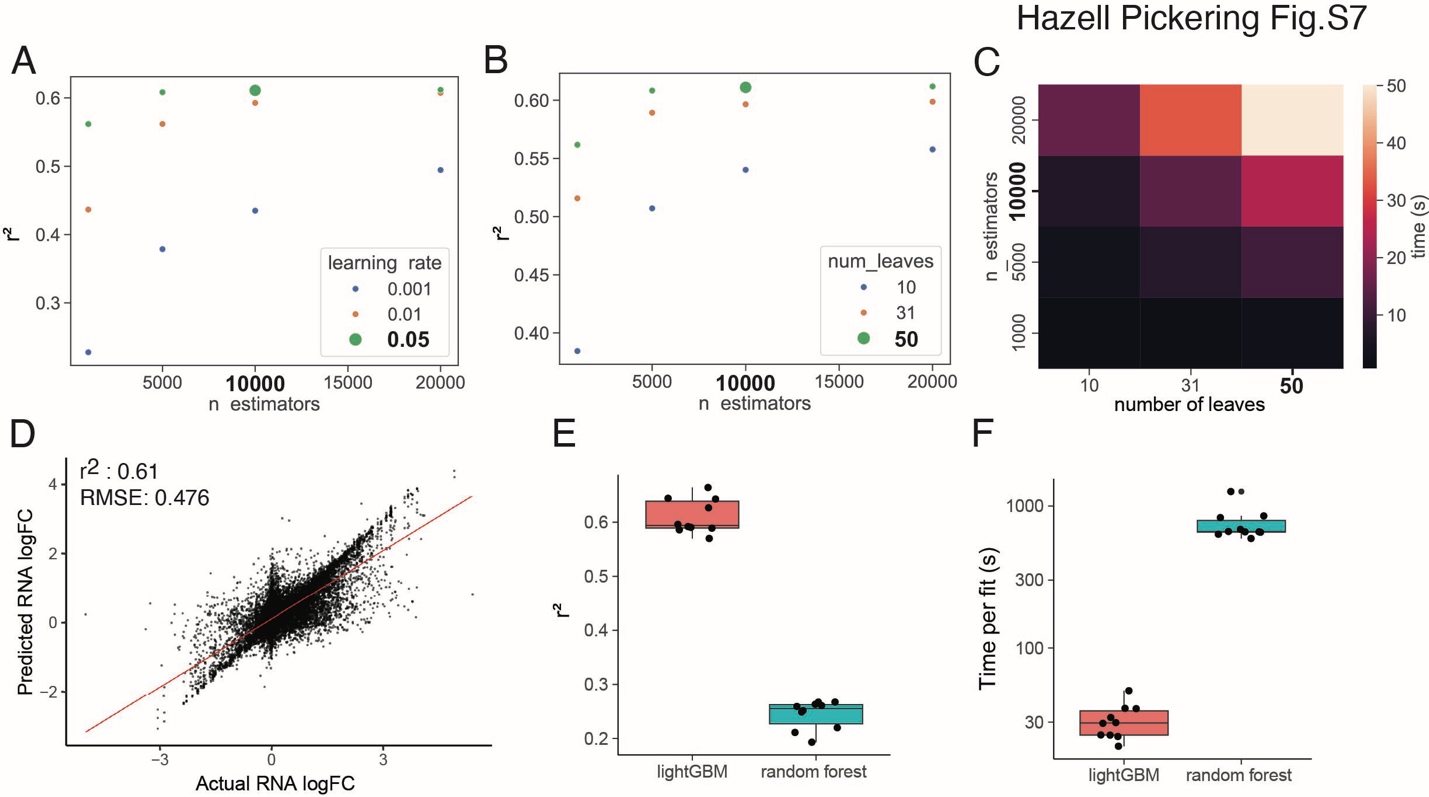

**Fig. S7.** Related to Fig3. **A** Average model performance (r^2^) for lightGBM models with different numbers of estimators and learning rates or **B** maximum number of leaves. Selected hyperparameter values are highlighted. **C** Average time taken to fit lightGBM models with different numbers of estimators and maximum number of leaves. **D** Scatter plot of actual vs predicted RNA log2 fold change of model trained with the best hyperparameters with a linear trendline. **E** Comparison of lightGBM and random forest (k=10) model performance and **F** time taken to fit. RMSE: root mean squared error.

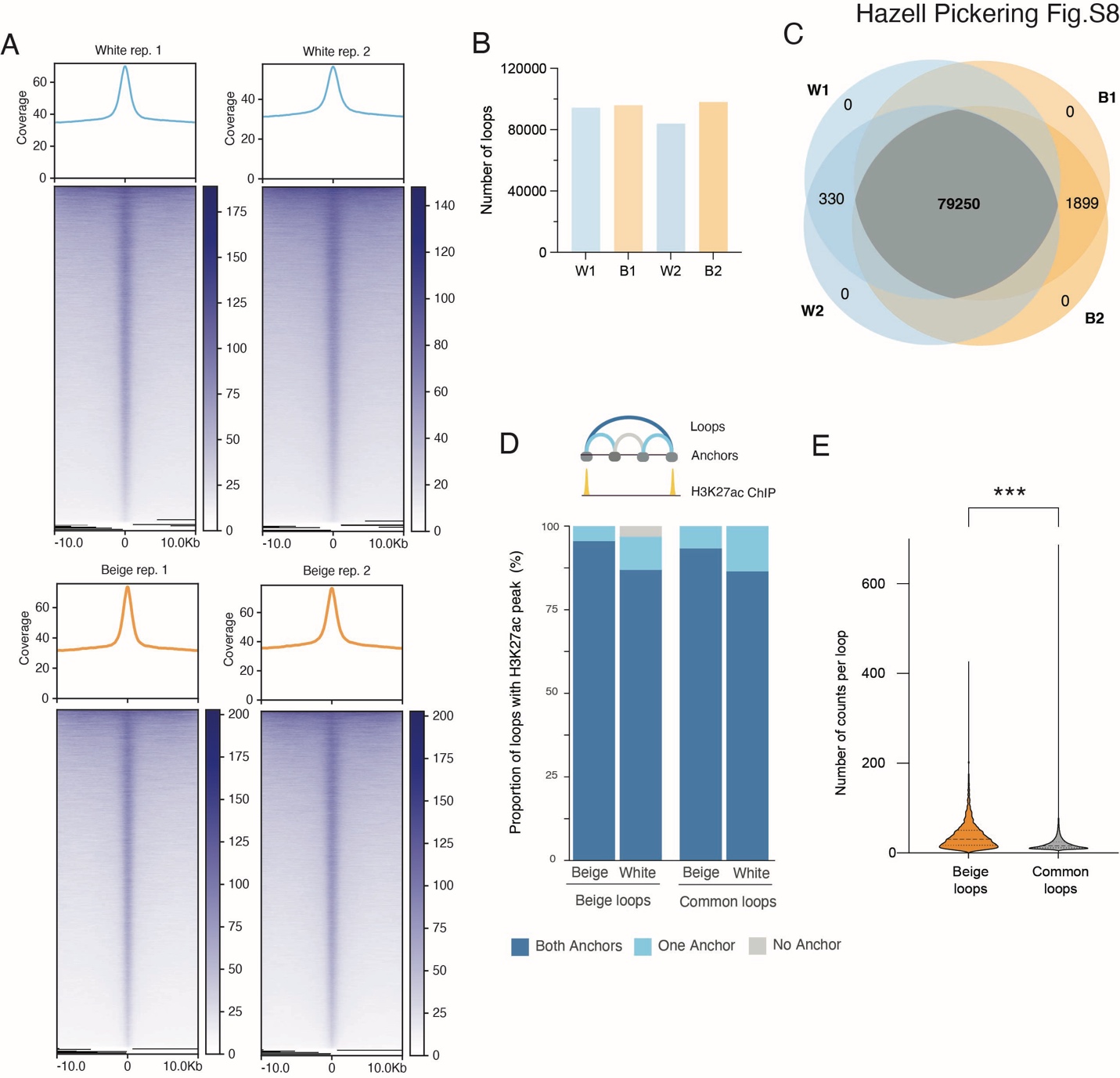

**Fig. S8**. Related to Fig4. **A** Hi-ChIP signal enrichment at ChIP peaks per condition (White/Beige) and per replicate. **B** Number of called loops per replicate (≥ 6 counts). **C** Euler diagram showing overlap of called loops for each replicate. **D** Proportion of loop anchors overlapping with white or beige H3K27ac peaks from an independent H3K27ac ChIP-seq experiment. **E** Number of counts per loop in beige and common loops (***p < 0.0001, two-tailed Mann-Whitney test; dashed lines represent the median and quartiles).

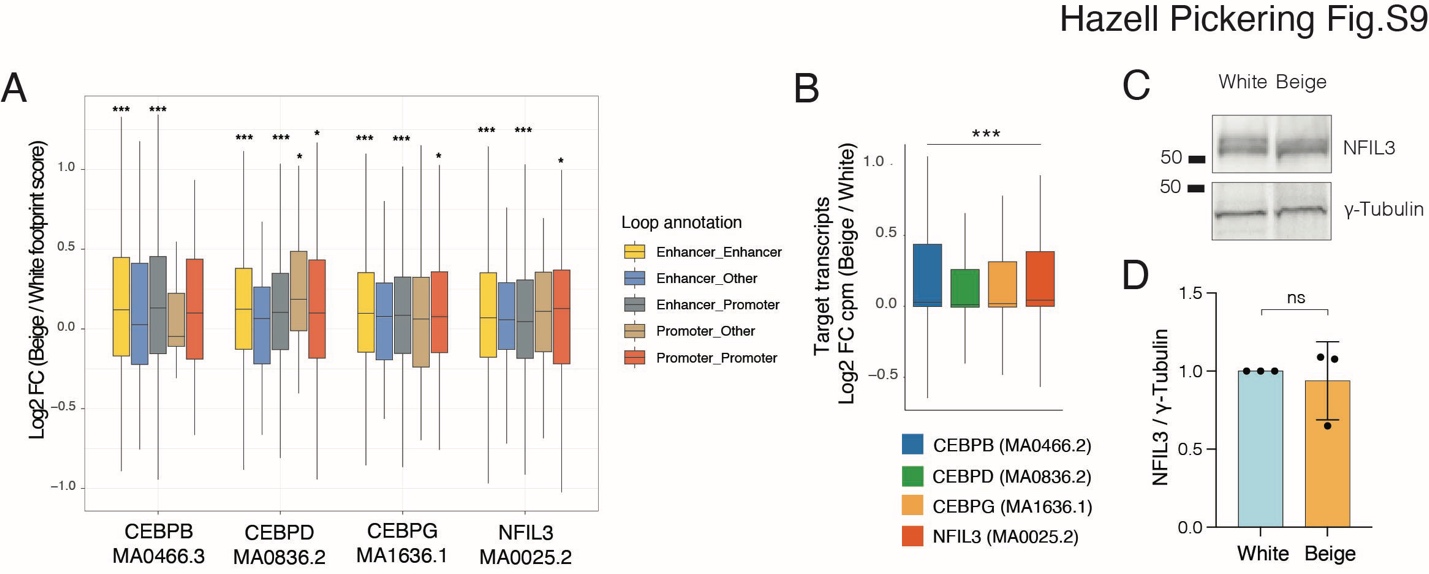

**Fig. S9.** Related to Fig5. **A** Log2 fold change footprint scores per loop category for the indicated TF motifs. **B** Log2 fold change beige vs white expression (normalized CPM over 300 bp downstream TSS) of transcripts with detected binding of indicated TFs at the promoter region. **C** Western blot analysis and **D** quantification of NFIL3 expression in white and beige adipocytes (D15) (mean fold difference ± SD; ns non-significant, Wilcoxon signed ranked test; n = 3). γ-Tubulin is shown as a loading control.

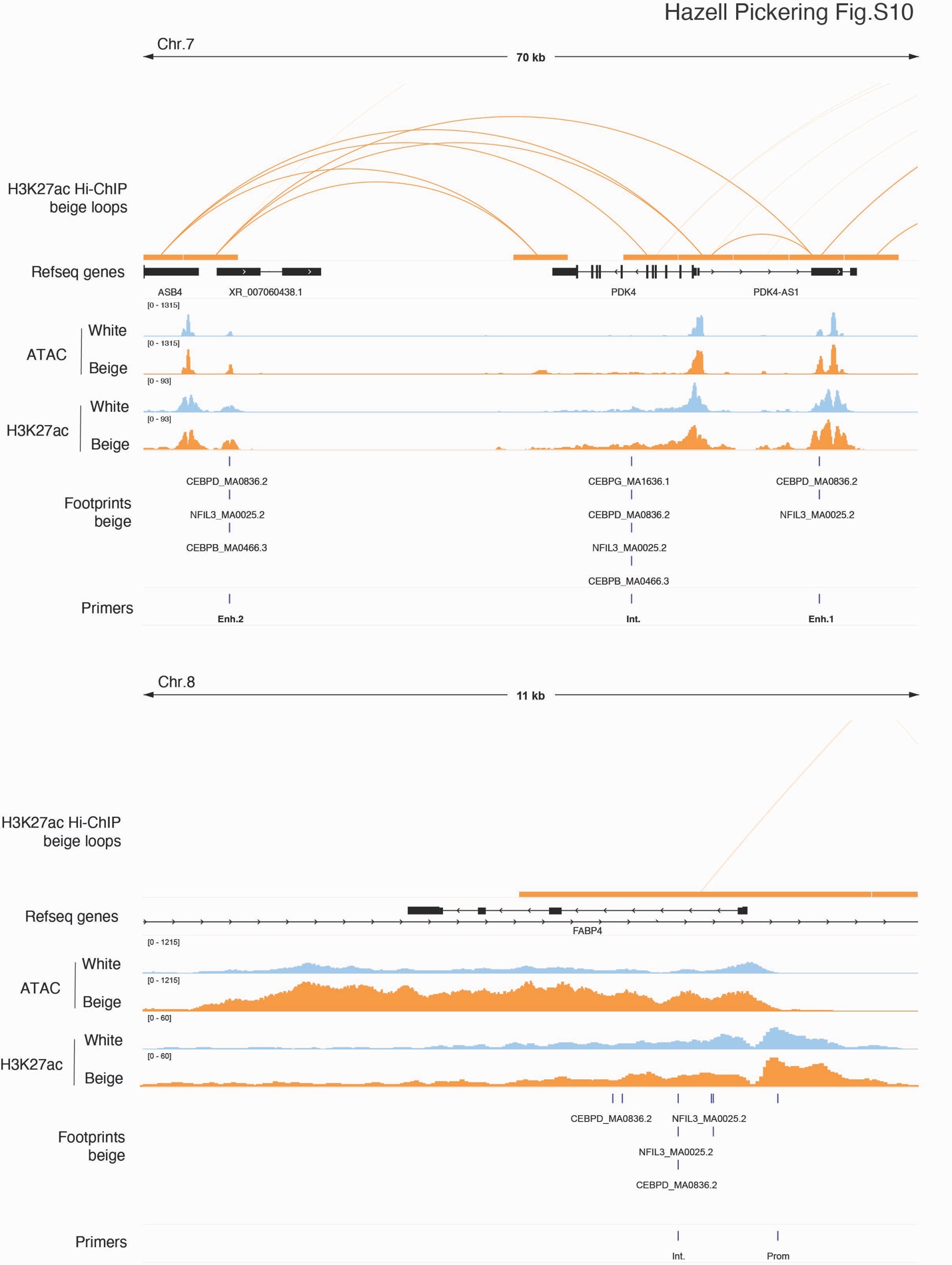

**Fig. S10.** Related to Fig5. Genome browser views of H3K27ac Hi-ChIP beige loops, ATAC, H3K27ac ChIP, C/EBP transcription factor binding sites identified by ATAC footprinting and primer location at PDK4 (top) and FABP4 (bottom) loci.
